# Cardiac acetylcholinesterase and butyrylcholinesterase have distinct localization and function

**DOI:** 10.1101/2024.05.29.596481

**Authors:** Dominika Dingová, Matej Kučera, Tibor Hodbod, Rodolphe Fischmeister, Eric Krejci, Anna Hrabovská

## Abstract

Cholinesterase (ChE) inhibitors are under consideration to be used in the treatment of cardiovascular pathologies. A prerequisite to advancing ChE inhibitors into the clinic is their thorough characterization in the heart. The aim here was to provide a detailed analysis of cardiac ChE to understand their molecular composition, localization, and physiological functions. A battery of biochemical, microscopic, and physiological experiments was used to analyze two known ChE, acetylcholinesterase (AChE) and butyrylcholinesterase (BChE), in hearts of mutant mice lacking different ChE molecular forms. Overall, AChE activity was exceeded by BChE, while it was localized mainly in the atria and the ventricular epicardium of the heart base. AChE was anchored by ColQ in the basal lamina or by PRiMA at the plasma membrane and co-localized with the neuronal marker TUJ1. In absence of anchored AChE, heart rate was unresponsive to a ChE inhibitor. BChE, the major ChE in heart, was detected predominantly in ventricles, presumably as a precursor (soluble monomers/dimers). Mice lacking BChE were more sensitive to a ChE inhibitor. Nevertheless, the overall impact on heart physiology was subtle, showing mainly a role in cholinergic antagonism to the positive inotropic effect of β-adrenergic stimulation. Our results help to unravel the mechanisms of ChE in cardiovascular pathologies and provide a foundation to facilitate the design of a novel, more effective pharmacotherapies, which may reduce morbidity and mortality of patients with various heart diseases.

**Abstract figure legend:** 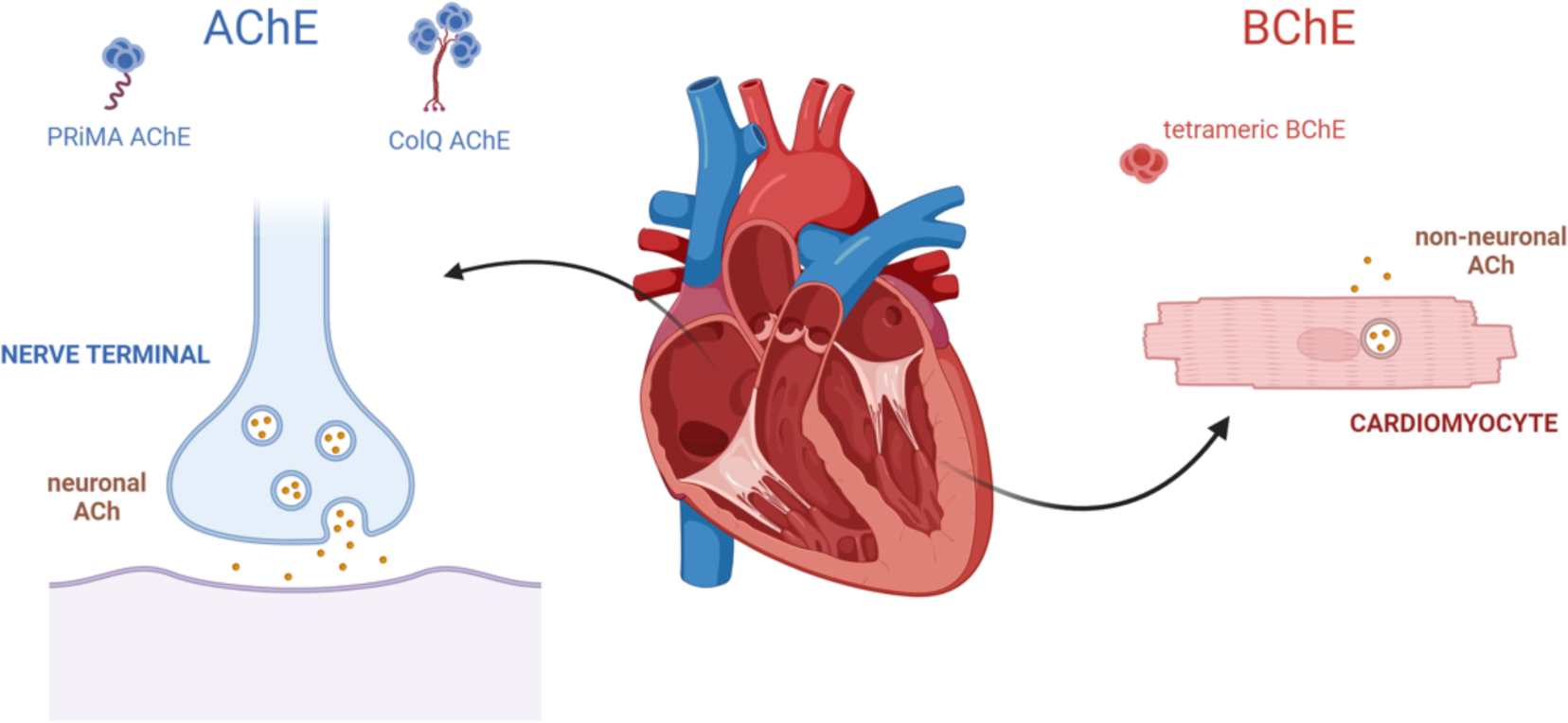

Acetylcholinesterase (AChE) has the highest activity in the atria. It is present in the heart in molecular forms anchored by a proline-rich membrane anchor (PRiMA) and by collagen Q (ColQ) and hydrolyzes acetylcholine of neuronal origin (neuronal ACh). Butyrylcholinesterase (BChE) is predominant in the ventricles. It is secreted in the form of a soluble tetramer and hydrolyzes acetylcholine originating from cardiomyocytes (non-neuronal ACh).

**Key points:**

- Inhibition of cholinesterases has therapeutic potential in cardiovascular pathologies
- Both known cholinesterases are present in heart
- Each cholinesterase has distinct localization patterns in the heart and functions in cardiac physiology
- Selective inhibition of acetylcholinesterase or butyrylcholinesterase may be used to alter specific cardiac functions
- Butyrylcholinesterase polymorphism may have an impact on the outcome of the cholinesterase inhibitor treatment

## Introduction

Cholinesterase (ChE) inhibitors (ChEIs) are clinically used in multiple indications, but mainly to treat cognitive deficits in patients with Alzheimer’s disease (Giacobini, 2004) and muscle weakness in patients with myasthenia gravis (Maggi & Mantegazza, 2011). By blocking degradation of acetylcholine (ACh), their therapeutic action is attributed to enhanced cholinergic transmission at synapses in the central nervous system and in the periphery (Hrabovska & Krejci, 2014). Recently, beneficial effects of ChEIs on the cardiovascular system of treated patients has been shown (Hsieh Mj *et al*., 2022). Indeed, the positive effects of increased cholinergic tone on cardiovascular functions has been confirmed in animal models and in patients with various cardiovascular diseases such as heart failure, arrhythmia, ischemia/reperfusion injury and hypertension (Rocha-Resende C *et al*., 2021). ChEIs decrease heart rate, beat to beat fluctuations, excitability, contractility, increase heart rate recovery after exercise, prevent cardiac remodeling, modulate angiogenesis and induce anti-inflammatory cell recruitment (Rocha-Resende C *et al*., 2021). These broad actions of ChEIs on heart function prompted the idea of developing large clinical trials with ChEIs in heart failure (Roy *et al*., 2015). However, a prerequisite to a large clinical study with ChEIs is to provide a detailed characterization of ChEs in heart and to unravel their role in cardiac physiology.

Two ChEs have been characterized in the body, acetylcholinesterase (AChE) and butyrylcholinesterase (BChE), which differ in their efficiency in hydrolysing ACh (Nicolet Y *et al*., 2003). AChE and BChE are organized by proline-rich association domains (PRAD) of different origin (Lockridge & Schopfer, 2020) into various molecular forms that differ in their oligomerization and localization (Massoulié, 2002). In the absence of PRAD, AChE and BChE monomers are likely retained in the endoplasmic reticulum where tetramers are normally assembled by PRAD (Dobbertin et al., 2009; Petrov et al., 2014). Collagen Q (ColQ) organizes AChE into tetramers and clusters AChE at the NMJ where AChE is critical to terminate the synaptic transmission. More than 30 mutations in ColQ in mouse or human are responsible of a congenital myasthenic syndrome, a severe muscular disease (Mihaylova *et al*., 2008; Wargon *et al*., 2012). The proline-rich membrane anchor (PRiMA) organizes AChE or BChE into tetramers and anchors these tetramers to the plasma membrane (Perrier *et al*., 2002). PRiMA anchors 90% of AChE activity in brain. In absence of PRiMA, mice adapt remarkably to the excess of ACh (Farar *et al*., 2012).

Both AChE and BChE are expressed in the heart, with BChE activity exceeding that of AChE (Gómez et al., 1999; Li et al., 2000). The main ChE anchoring proteins, i.e., PRiMA and ColQ, are expressed in the heart at the mRNA level (Perrier *et al*., 2002; Kilianova *et al*., 2020). Assembly of AChE and BChE with ColQ has been confirmed in rat ventricle (Krejci *et al*., 1997).

Despite a clear role of ChEs in ACh hydrolysis, their properties have yet to be characterized in the heart. Cardiac ACh originates from at least two sources: *nervus vagus,* being part of the cardiac neuronal cholinergic system, and cardiomyocytes, as part of the cardiac non-neuronal cholinergic system (Rocha-Resende C *et al*., 2021). These two systems work in synergy to counterbalance cardiac sympathetic stimulation (Roy *et al*., 2015), but with different physiological and pathophysiological properties.

The aim of this work was to provide a detailed analysis of cardiac ChEs and to understand their physiological functions. We characterized the ChEs in mouse heart by assessing their activity, localization of functional forms, and role in cardiac physiology. By utilizing mutant mouse strains lacking ChE or their anchoring proteins (Feng *et al*., 1999; Camp *et al*., 2005; Li *et al*., 2008; Dobbertin *et al*., 2009), we also investigated the potential impact of their selective absence or inhibition in specific disorders.

## Materials and methods

### Experimental animals

All experiments were performed following French and Slovak guidelines for laboratory animal handling. The protocols were approved by the Ethical Committee of the Faculty of Pharmacy of Comenius University in Bratislava, the State Veterinary and Food Administration of the Slovak Republic (Ro-557/11-221/2, Ro-1786/12-221), the Ethical Committee of Université Paris-Saclay, and the Ethical Committee of Université Paris Descartes, in accordance with the European Community Council Directive of February 1, 2013 (82010/63/EEC registration number CEEA34.EK/AGC/LB.111.12).

Adult male and female WT mice and mutant mouse strains were used in the experiments. Specifically, we used BChE^−/−^ mice (Li *et al*., 2008), with a complete absence of BChE, AChE del E 5+6^−/−^ mice (Camp *et al*., 2005 p.2005) which lack all anchored AChE forms thus producing only soluble AChE monomers; ColQ^−/−^ mice (Feng *et al*., 1999), which do not produce basal lamina anchored AChE or BChE tetramers; and PRiMA^−/−^ mice (Dobbertin *et al*., 2009), which lack PRiMA protein and consequently do not anchor AChE or BChE to the plasma membrane. All experimental animals were of mixed genetic background equivalent to F2 and F3 generation of mating of C57B6 and DBA2 strains to exclude strain specific properties. The age of the mice varied from 3-15 months, an age frame that has little impact on cardiac function (Dai *et al*., 2012), neuronal and neuromuscular nicotinic receptor expression, or neuronal ACh biosynthesis and degradation (Sarter & Bruno, 1998).

### Heart isolation

Mice were decapitated and the beating hearts were explanted in chilled Krebs-Henseleit solution (117.9 mM NaCl, 4.7 mM KCl, 1.2 mM MgSO_4_ x 7 H_2_O (Sigma Aldrich), 1.2 mM KH_2_PO_4_, 0.4 mM CaCl_2_, 25.04 mM NaHCO_3_, and 15.5 mM glucose) and cannulated through the aorta. Hearts were then perfused following Langendorff’s protocol (Georget *et al*., 2003) for 20 min under constant pressure (75 mmHg) with Krebs-Henseleit solution (117.9 mM NaCl, 4.7 mM KCl, 1.2 mM MgSO_4_ x 7 H_2_O (Sigma Aldrich), 1.2 mM KH_2_PO_4_, 1.9 mM CaCl_2_, 25.04 mM NaHCO_3_, 11 mM glucose, 0.1 mM EDTA, 2 mM sodium pyruvate (Bio Basic Canada), and 1.1 mM mannitol (Bio Basic Canada)). The solutions were constantly stirred and oxygenated with 95% O_2_ and 5% CO_2_ at 37 ± 0.2°C. Unless stated otherwise, all chemicals were purchased from Euromedex.

### Biochemical Analysis

After Langendorff perfusion, whole hearts were used or hearts were immediately dissected into either the right and left atria, right and left ventricles, and septum or into the base and the apex of the heart, flash frozen in liquid nitrogen and stored at −80°C until further analysis.

### AChE and BChE expression

The mRNA was extracted from frozen tissues using a standard phenol/chloroform extraction method with TRI-Reagent (Sigma-Aldrich). Tissues were homogenized with steel beads using the TissueLyser II (Qiagen). The concentration of isolated mRNA was determined utilizing the Take3 Microvolume plate (BioTek) and Synergy H4 Hybrid Reader (BioTek). Reverse transcription was carried out in a T100™ Thermal Cycler (Bio-Rad) employing the High-Capacity cDNA Reverse Transcription Kit (Applied Biosystems), following the manufacturer’s protocol. RT-qPCR analysis was conducted using SYBR Select Master Mix (Applied Biosystems) on the Quant Studio 3 PCR system (Applied Biosystems). The expression levels of AChE (Forward: 5’-TTTTCCTTCGTGCCTGTGGT; Reverse: 5’-GGGACCCCGTAAACCAGAAA) and BChE (Forward: 5’-AACTTCGTGCTCCCCAATGT; Reverse: 5’-ATCCTGCCTTCCACTCTTGC) were normalized to that of the housekeeping gene *Actb* (Forward: 5’-CGGCTTTGCACATGCCGGAGC; Reverse: 5’-GTCCACACCCGCCACCAGTTCG).

### Tissue extract preparation

Frozen tissue was transferred into pre-chilled 2 ml microfuge tubes containing two 5 mm stainless steel beads (Qiagen) and 5 volumes of ice-cold buffer (10 mM HEPES buffer pH 7.5, 10 mM EDTA, 0.8 M NaCl, and 1% CHAPS (Euromedex)) and agitated for 2.5 min at a frequency of 25 Hz in a Mixer Mill MM 300 (Retsch). Homogenates were then centrifuged at 14000 g for 10 min at 4°C and supernatants representing the tissue extracts were transferred into clean pre-chilled microcentrifuge tubes and used immediately in experiments. Proteins were quantified using commercially available Pierce BCA Protein Assay Kit (Thermo Scientific).

### ChE activity

ChE activity was measured in tissue extracts by modified Ellman’s assay (Dingova *et al*., 2014). In AChE activity measurements, BChE activity was inhibited by 30 min incubation with 20 μM tetra(monoisopropyl)pyrophosphortetramide (iso-OMPA, Sigma Aldrich) prior to incubation with the substrate, 1 mM acetylthiocholine (ATC, Sigma Aldrich). BChE activity was determined with 1 mM butyrylthiocholine (BTC, Sigma Aldrich). The reaction was run in 5 mM HEPES buffer pH 7.5 in the presence of 0.5 mM 5,5’-dithiobis-(2-nitrobenzoic acid) and monitored at 415 nm for 30 min at 25°C.

### Molecular forms of ChE in heart

Molecular forms of ChE in extracts were separated in a 5-20% sucrose gradient (Bernard *et al*., 2011) containing 10 mM HEPES buffer pH 7.5, 0.8 M NaCl, 10 mM EDTA and 0.2% BRIJ (Sigma Aldrich) or 1% CHAPS. β-galactosidase (Sigma Aldrich) and alkaline phosphatase (Sigma Aldrich) were used as sedimentation standards. Gradients were centrifuged in a pre-chilled rotor SW41 at 38000 rpm for 17 hours at 7°C in a Beckman Coulter Optima LE-80K. Approximately 48 fractions were collected by an electric pump and distributed into a clean 96-well plate for further analysis. AChE and BChE activities were determined by modified Ellman’s assay as described above (Dingova *et al*., 2014). The area under the curve (AUC) was calculated using GraphPad Prism 8.4.3 (GraphPad Software, San Diego). Activity of β-galactosidase was determined using 3.15% 2-nitrophenyl β-D-galactopyranoside (Sigma Aldrich) in 0.1 M HEPES buffer pH 7.5, 0.1 M NaCl, and 0.01 M MgCl_2_ (Sigma Aldrich) at 415 nm. Activity of alkaline phosphatase was detected in the presence of 1.35% p-nitrophenyl phosphate (Sigma Aldrich) in 0.1 M TRIS buffer pH 8.0 and 0.01 M MgCl_2_ (Sigma Aldrich) at 415 nm.

The profile of AChE and BChE activity of heart extract separated in sucrose gradient is highly dependent on the perfusion. When heart is not perfused, a peak of AChE activity dominates at 4S and contains dimers anchored by glycosylphosphatidylinositol on red blood cells; a peak of BChE activity dominates at 12S and contains soluble BChE tetramer. Indeed, we observed correlated variations of AChE 4S peak and BChE 12S peak that presumably reveal variation in perfusion. The perfusion is thus a critical step to quantify ChE in highly vascularized tissues such as heart.

### Microscopic Analysis

#### Cryosections

After Langendorff perfusion, hearts were prepared for microscopic studies by immediate immersion into 3 M KCl to stop its function in diastole and placed into 20% sucrose (Sigma Aldrich) for 2 hours at 4°C, followed by 40% sucrose for 2 hours at 4°C. Hearts were then placed into molds, embedded in O.C.T. (Tissue-Tek, Sakura) and slowly frozen in isopentane (Sigma-Aldrich) cooled with dry-ice to −30°C. Tissues were cut to 12 µm thick transverse or longitudinal sections in a cryostat, mounted on SuperFrost slides (Fisher), and stored at −80°C. Slides were defrosted before use, fixed with 4% paraformaldehyde (Sigma Aldrich) for 15 min at room temperature and washed in distilled water.

### Whole-mounted hearts

Once the hearts were stopped in diastole, warm 20% liquid gelatin was injected into all four heart chambers to inflate its volume (Pauza *et al*., 2000). Subsequently, hearts were placed into the ice-cold 0.01 M phosphate buffer, pH 7,4 containing 0,15 mM NaCl (PBS), allowing the gelatin to polymerize, and thus keeping the hearts inflated (Fig. 1*B*). Gelatin-filled hearts were rinsed in PBS and fixed with 4% paraformaldehyde for 30 min at 4°C. For better penetration of the staining solution, hearts were incubated with hyaluronidase (0.5 mg/100 ml, Sigma Aldrich) in 0.1 M PBS overnight at 4°C. Tissues were washed briefly in PBS the next day and used for further analysis.

**Figure 1.**
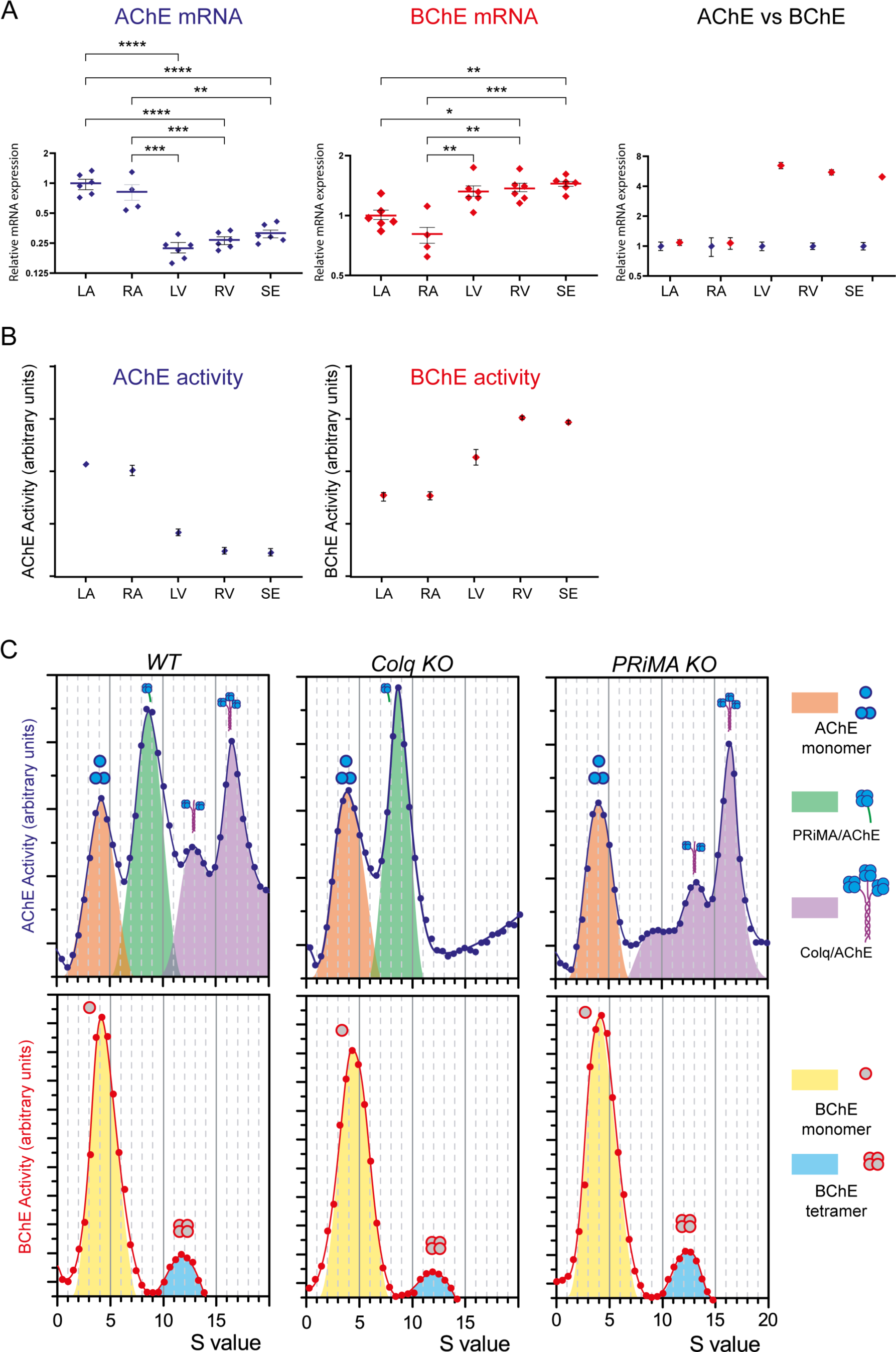
Distribution of ChE activity in Langendorff-perfused mouse heart compartments. ***A*** Expression of AChE and BChE in left atrium (LA), right atrium (RA), right ventricle (RV), left ventricle (LV) and septum (SE) in reference to LA (*left* for AChE; *middle* for BChE) and in reference to AChE in each heart compartment (right). AChE showed higher expression in atria than in ventricles (*p ≤ 0.0001)*, while BChE expression predominated in ventricles (*p ≤ 0.0001)*. Both ChEs were expressed comparably in atria, while expression of BChE exceeded AChE in ventricles (*p ≤ 0.0001)*. Values are presented as mean ± SEM, p ≤ 0.05 (*), p ≤ 0.01 (**), p ≤ 0.001 (***), p ≤ 0.0001 (****), n=6 with exception for RA where n=4 due to insufficient RNA. ***B*** Activity pattern of AChE and BChE in left atrium (LA), right atrium (RA), right ventricle (RV), left ventricle (LV) and septum (SE) measured in a pool of four mice, ran in duplicate. AChE activity predominated in atria (*p ≤ 0.0001)* and BChE activity in ventricle (*p ≤ 0.0001).* AChE exceeded BChE activity in atria (left atria: p = 0.00051, right atria: p = 0.0061), while BChE activity dominated in the ventricles (*p ≤ 0.0001*). Values are presented as mean ± SEM, p ≤ 0.05 (*), p ≤ 0.01 (**), p ≤ 0.001 (***), p ≤ 0.0001 (****). Experiments were repeated at least three times. **C** Molecular forms assembled in perfused murine hearts. Proteins were extracted with detergent and high salt (0.8M NaCl) to solubilize anchored forms from membrane and basal lamina and separated in sucrose gradient containing BRIJ and high salts. Each graph represents ChE activity in each collected sucrose gradient fraction (blue dots for AChE, red dots for BChE). AChE activity (top panels) distributes in four major peaks. The red peak at 4S corresponds to monomer and dimer of AChE, the green peak at 9S corresponds to PRiMA AChE complex, the purple peaks at 12.5S and 16S to ColQ AChE complex. BChE activity (lower panels) distributes in two peaks. The yellow peak at 4.5S corresponds to monomer of BChE and the blue peak at 12S corresponds to BChE tetramer.

### Tsuji’s ChE activity staining method

Slides with mounted cryosections were pre-incubated for 30 min with selective inhibitors of ChE. BW284C51 (BW, 1 µM, Sigma Aldrich) was used to inhibit AChE and/or iso-OMPA (20 µM, Sigma Aldrich) to inhibit BChE. Subsequently, sections were stained for AChE or BChE activity by modified Karnovsky and Roots’ staining procedure (Tsuji *et al*., 2002). Briefly, slides were incubated at room temperature for 6 hours in staining solution containing 15 mM citrate buffer pH 5.3 (Sigma Aldrich), 2.8 mM CuSO (Sigma Aldrich), 0.47 mM K_3_FeCN_6_ (Sigma Aldrich), and 1.7 mM substrate. AChE activity was revealed using substrate ATC in the presence of iso-OMPA (20 µM). BChE activity was visualized with substrate BTC. Staining solution for negative control contained BW (1 µM) and iso-OMPA (20 µM). Slides were then washed in 0.01 M citrate buffer, pH 5.3 at room temperature, gradually dehydrated in ethanol (50-100%), immersed in xylene substitutes Ottix (DIAPATH Microstain division) for 10 min at room temperature and mounted with Eukitt medium (Sigma-Aldrich). Activity was visualized by light microscopy (Olympus BX61) and pictures were processed in Adobe Photoshop or GIMP.

Activity staining of gelatin-filled heart followed the same procedure, but the incubation was shortened to 3 hours at room temperature. Subsequently, hearts were properly washed and post-fixed in 4% paraformaldehyde. Tissues were then placed into a Petri dish filled with distilled water and observed using light microscopy (Olympus BX61). As these preparations were 3D objects, multiple pictures with consecutive planar focuses were taken for the final pictures, which were composed using extended depth of field in ImageJ program (Hein *et al*., 2012) and processed in Adobe Photoshop or GIMP.

### Immunohistochemistry

Transverse and longitudinal sections of heart were traced with a Dako pen (Dako, S2002). Slides were covered with 50 mM glycine for 10 min and subsequently washed with PBS. Dehydration of slides was performed by graded ethanol 50-100% (each grade for 5 min). For better penetration of antibodies, Dent’s solution (DMSO:CH_3_OH in a 1:4 ratio) was applied for 15 min. To enhance fluorescence contrast, 30% H_2_O_2_:Dent’s solution in a ratio 1:4 was applied for 1 hour. Samples were rehydrated in graded ethanol (100-50%) and washed 3 times for 5 min in PBS. Non-specific sites were blocked with 5% goat serum diluted in PBS for 30 min and subsequently washed 3 times for 5 min with PBS. Sections were incubated for 2 hours with rabbit polyclonal antibody against mouse AChE (dilution 1:500), which was a gift from Prof. Palmer Taylor (University of California, San Diego, La Jolla, CA, USA). Slides were washed with PBS 5 times for 5 min. Subsequently, DyLight 594 goat anti-rabbit IgG (dilution 1:500, Vector Laboratories DI-1594) secondary antibody was applied for 30 min and thoroughly washed with PBS for 1 hour (the solution was changed every 10 min). Slides were then mounted with Fluoroshield (Sigma-Aldrich) or Fluoroshield with DAPI (Sigma-Aldrich). All steps were performed at room temperature and protected from light. For co-localization of AChE protein with a neuronal marker, a mouse monoclonal antibody against mouse neuronal class III β-tubulin (TUJ1) labelled with Alexa Flour 488 (dilution 1:500, Covance, A488-435L) was incubated together with primary antibody against mouse AChE. Immunofluorescence was observed with fluorescence microscopy (Nikon Eclipse Ni-U) and pictures were processed in Adobe Photoshop (2014) or GIMP (2020).

### Left ventricular catheterization

Animals were anesthetized intraperitoneally by 2.5% solution of 2,2,2-tribromoethanol - AVERTIN (Sigma Aldrich, USA, dose 15 µl/g) and placed on a 37°C heating table. The depth of anaesthesia was confirmed by pedal reflex (firm toe pinch). After a 20-min stabilization phase, basal hemodynamic parameters were recorded in the left ventricle following published protocols (Müller *et al*., 2003; Lewin *et al*., 2009). In short, a polyethylene catheter (0.62 mm, Smith Medical International Ltd, Great Britain), connected to pressure detector ELS–01 (Experimentia LTD, Hungary), was filled with heparinized physiological solution (63 UI/ml, Zentiva a.s, Czech Republic) and inserted via right carotid artery into the left ventricle and left ventricular pressure curve was recorded. Using the S.P.E.L. Advanced Haemosys program (SOFT-01-HAEMO, Experimentia Ltd., Hungary), heart rate, left ventricular systolic pressure (LVSP), diastolic pressure (LVDP), and maximum rate of ventricular contraction (dP/dt_max_) and relaxation (dP/dt_min_) were obtained. Finally, the β_1_-adrenoreceptor agonist dobutamine (dobutamine hydrochloride, Sigma Aldrich, USA) was cumulatively injected (every 3 min for 12 min) via the cannulated left jugular vein by automated infusion pump. The absolute cumulative dose of dobutamine was 166.80 µg/kg. While still under anesthesia, animals were euthanized at the end of the experiment by cervical dislocation. Experiments were performed with female WT mice, PRiMA^−/−^ and BChE^−/−^ mice. ColQ^−/−^ and AChE del E 5+6^−/−^ mice, which are characterized by their smaller size (Feng *et al*., 1999; Camp *et al*., 2005; Dobbertin *et al*., 2009), had smaller venous lumen size and increased fragility of the *arteria carotis*, and thus hemodynamic experiments were not possible to perform.

### Isolated perfused heart assay

#### Spontaneously beating hearts

Hearts were explanted, cannulated, and mounted on Langendorff’s apparatus as described in the paragraph *Heart isolation*. Subsequently, a small latex water-filled balloon was inserted into the left ventricle and connected through a catheter to a pressure transducer ELS-01 (Experimentia Ltd, Hungary) with pressure set to 5-12 mmHg. A left ventricular pressure curve was recorded and subsequently heart rate was measured. The analysis was performed with S.P.E.L Advanced Heamsys software (SOFT-01-HAEMO, Experimentia Ltd, Hungary). All studied substances were applied by an infusion pump SN 50F6 (Sino Medical-Device Technology, Co., Ltd., China), while the application rate was synchronized to the actual coronary blood flow, unless stated otherwise. After a 20 min stabilization phase, basal physiological parameters were recorded for the following 5 min. The effect of endogenous ACh on heart rate was examined during an application of the non-selective inhibitor of ChE, neostigmine (3 µM, Sigma Aldrich, USA), for 25 min. Involvement of the muscarinic or nicotinic receptors was studied using the muscarinic antagonist, atropine (1 µM, Sigma Aldrich, USA), or the nicotinic antagonist, hexamethonium (0.1 mM, Sigma Aldrich, USA). After 10 min perfusion with 3 µM neostigmine, either atropine or hexamethonium were added to the perfusion buffer and heart rate was recorded for an additional 10 min. The effect of exogenous ACh on the heart was studied under β-adrenergic receptor stimulation by isoproterenol (ISO, (−)- isoproterenol, Sigma Aldrich, USA). A progressively increasing dose of ISO (0.1 nM, 0.3 nM, 1 nM, 3 nM, 10 nM, 30 nM, 100 nM) was applied in 7 min intervals. During application of the highest concentration of ISO, exogenous ACh was added in concentrations of 0.3 µM, 1 µM and 10 µM. Finally, the effect of ACh was blocked by atropine (0.1 µM). Experiments were performed with male WT mice, PRiMA^−/−^ and BChE^−/−^ mice. ColQ^−/−^ and AChE del E 5+6^−/−^ mice, which are characterized by their smaller size (Feng *et al*., 1999; Camp *et al*., 2005; Dobbertin *et al*., 2009), had smaller venous lumen size and increased fragility of the *arteria carotis*, and thus physiological experiments were not possible to perform.

### Paced hearts

Hearts were explanted, cannulated, and mounted on Langendorff’s apparatus under the same conditions as described in the chapter *Heart isolation and tissue processing*. The experiment was performed under paced conditions (650 bpm) via platinum wires at reduced calcium concentration (1.1 mM). The balloon was connected to a pressure transducer (Statham gauge Ohmeda, Bilthoven, Netherlands) and the pressure was set to the conditions of isovolumic contraction. IOX2 (Emka technologies, Paris, France) was used to obtain the values of left ventricular developed pressure (LVDevP), end-diastolic pressure (EDP), dP/dt_max_ and dP/dt_min_ (Transonic Systems T106, Transonic Systems Inc., USA). Basal values were recorded after a 10 min equilibration period. Subsequently, drugs were applied into a perfusion buffer by an infusion pump (Pump 11, Harvard Apparatus, USA). Cholinergic stimulation of heart by endogenous ACh was achieved by application of neostigmine (3 µM) for 15 min. Stimulation of β-adrenergic receptors was accomplished by infusion of progressively increasing concentrations of ISO (0.1, 0.3, 1, 3, 10, 30 and 100 nM) in 7 min intervals. During application of the highest ISO concentration, ACh (0.3 µM), neostigmine (3 µM) and ACh (10 µM) were applied in 10 min intervals. Finally, the effect of ACh on muscarinic receptors was blocked by atropine (0.1 µM). Experiments were performed with male WT mice, PRiMA^−/−^ and BChE^−/−^ mice. ColQ^−/−^ and AChE del E 5+6^−/−^ mice, which are characterized by their smaller size (Feng *et al*., 1999; Camp *et al*., 2005; Dobbertin *et al*., 2009), had smaller venous lumen size and increased fragility of the *arteria carotis*, and thus physiological experiments were not possible to perform.

### Data analysis

Data were processed in Microsoft Office Excel and GraphPad Prism 8.4.3 (GraphPad Software, San Diego). Values are expressed as mean ± SD. Statistically significant differences were determined by using one-way ANOVA followed by Bonferroni’s multiple comparison test or two-way ANOVA followed by Tukey or Dunnet’s post hoc test with p ≤ 0.05 (*), p ≤ 0.01 (**), p ≤ 0.001 (***), p ≤ 0.0001 (****).

## Results

### Biochemical analysis of cholinesterases in heart

Quantification of AChE and BChE in heart is complicated by the high levels of both enzymes in blood, AChE is abundant as a dimer anchored at the surface of red blood cells and BChE is abundant as soluble tetramer. To analyse the distribution of ChE in the heart, the beating hearts were therefore explanted in chilled Krebs-Henseleit solution and perfused by Langendorff’s method prior dissection to the compartments.

RT-qPCR analysis confirmed expression of both ChE in murine heart, with different expression patterns within the compartments (*Fig. 1A*). AChE was predominantly expressed in atria with expressions in ventricles being about 25% of the atrial values (*p ≤ 0.0001)*. In contrary, BChE expression was about 50% higher in ventricle than in atria (*p ≤ 0.0001).* Relative expressions of AChE and BChE were comparable in atria but AChE expression levels in ventricle represented only 15 – 20% of BChE values (*p ≤ 0.0001)*.

ChE activity was assessed using modified Ellman’s assay. Activity assay in the whole heart revealed lower AChE than BChE activity (AChE (ΔA/min = 0.0123), BChE (ΔA/min = 0.0282)). AChE exceeded BChE activity in atria (left atria: p = 0.00051, right atria: p = 0.0061; *Fig. 1B*), while BChE activity dominated in the ventricles (*Fig. 1B*; p ≤ 0.0001). AChE activity was comparable in both atria and about 2.5-fold lower in ventricles (*Fig. 1B*; p ≤ 0.0001). BChE activity was the highest in left ventricle and septum, followed by right ventricle (*Fig. 1B*; p ≤ 0.0001), and atria (*Fig. 1B*; p ≤ 0.0001).

To evaluate the relative contribution of different AChE and BChE complexes (molecular forms), we solubilized ChE from heart extract (see method) and separated the proteins in sucrose gradient. AChE activities of each fraction were distributed in four main peaks, a peak around 4S, a peak around 9S, a peak at 12.5S and a peak at 16S. The position of each peak was normalized with internal markers for standardization. AChE peak at 4S corresponds to monomer and dimer. AChE peak at 9S corresponds to PRiMA anchored tetramer (Perrier *et al*., 2002), that was confirmed by its absence in the PRiMA^−/−^ hearts. AChE peaks at 12.5S and 16S correspond to ColQ anchored tetramer, complex oligomers that associate a trimer of collagen with 2 or 3 tetramers of AChE respectively. As expected, these two peaks were absent in ColQ^−/−^ mice. Relative quantification of peaks by calculating area under the curve (AUC) of PRiMA AChE was 1.25-fold higher than ColQ AChE. Note that the peaks of ColQ AChE did not change in extract of PRiMA^−/−^ heart compared to the WT and the peaks of PRiMA AChE did not change in extract of ColQ^−/−^ heart compared to the WT.

In contrast to AChE, BChE activity was distributed in two peaks. The main peak at 4.5S corresponds to monomer of BChE. A minor peak at 12S corresponds to a soluble tetramer originating from blood. The profile of BChE is similar in extracts from WT, PRiMA^−/−^ and ColQ^−/−^ hearts thus BChE is not quantitatively anchored by PRiMA at the cell surface or by ColQ in the basal lamina.

Altogether, overall BChE activity was higher than AChE activity in the whole heart. Nevertheless, AChE was dominant in atria, where AChE is anchored by PRiMA at the plasma membrane and by ColQ in extracellular matrix. BChE activity was dominant in ventricles mainly as monomers. Monomer of BChE is the precursor of tetramer and is presumably retained in endoplasmic reticulum.

### Localization of cholinesterases in heart

AChE has been used as a marker to visualize neurons in atria, sinoatrial and atrioventricular nodes (Pauza *et al*., 2000). AChE is, however, quantitatively anchored either to the plasma membrane by PRiMA or to the extracellular matrix by ColQ, while we detected both these molecular anchored forms of AChE in the cardiac tissue in biochemical analyses described above. It therefore remains to know if the complexes have similar or different distribution in the heart. We thus took advantage of mutant mice to localize AChE anchored by ColQ (in PRiMA^−/−^ mice) and AChE anchored by PRiMA (in ColQ^−/−^ mice). Hearts were perfused to eliminate ChE originating from blood, stopped in diastole and inflated by injecting gelatin into all chambers to improve the surface visualization and stained for ChE activity. Staining by Tsuji’s method provided brown precipitate corresponding to ChE activity. Whole-mounted hearts were stained for ChE activity in epicardium. Cryosections were used to visualize ChE activity in myocardium and to colocalize the ChE protein with neuronal (TUJ1) marker by immunohistochemistry.

### Localization of cholinesterases in epicardium and myocardium of atria

Tsuji’s staining revealed strong, homogenously distributed ChE activity in epicardium (*Fig. 2 A)* and myocardium (*Fig. 2 B)*. This staining formed in fine branching was similar in heart of BChE^−/−^ mice, thus appertain to AChE staining. To confirm the specificity of the precipitate, we used hearts from AChE del E 5+6^−/−^ mice, in which the last exon of AChE gene is deleted. As consequence, AChE cannot interact with ColQ and PRiMA. The ChE staining was absent in atria of AChE del E 5+6^−/−^ mice. The staining was also absent in atria of PRiMA^−/−^ mice but not of ColQ^−/−^ mice where the staining was comparable to WT. Immunohistochemistry in WT transverse sections revealed anti-AChE antibody co-localized with the neuronal marker TUJ1 (Fig. 2*C*). Altogether, the AChE activity in atrial epicardium and myocardium depends on AChE anchored by PRiMA at the surface of neurons. This is a key function of PRiMA well established in the brain (Dobbertin *et al*., 2009). Despite ColQ AChE is abundant in extract, the deposition of AChE anchored by ColQ presumably in the basal lamina is not densely localized and is not detectable by the staining method.

**Figure 2.**
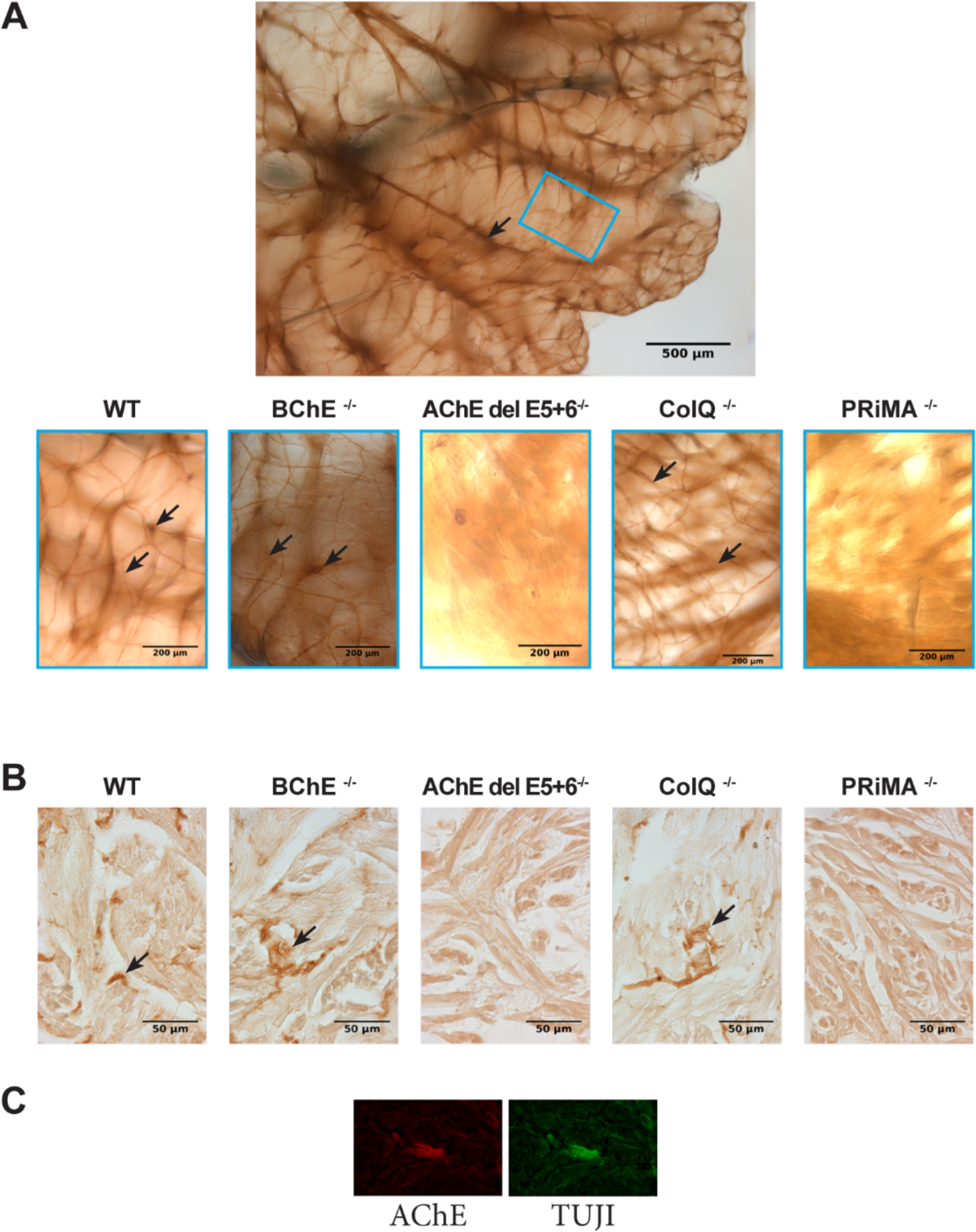
Localization of ChEs in atria. ***A,B*** Localization of ChE in atrial epicardium of the whole-mounted heart (***A***) and in cryosections (***B***) visualized by Tsuji’s staining for 3 hours with ATC. ChE activity is marked by black arrows. ChE activity was detected in WT, BChE^−/−^, and ColQ^−/−^ mice, but not AChE del E 5+6^−/−^ and PRiMA^−/−^ mice. ***C*** Co-localization of AChE and a neuronal marker (TUJ1) in the atria by immunohistochemistry.

### Localization of cholinesterase activities in ventricles

Staining by Tsuji’s method revealed strong, homogenously distributed ChE activity in the epicardium in the base of inflated WT hearts (*Fig. 3)* that gradually attenuated towards the apex. Similar results were acquired in BChE mutants. In AChE del E 5+6^−/−^ mice, only one markedly stained fiber-like structure (blue arrows in the Fig. 3), which disappeared after treatment with a BChE inhibitor, was consistently observed. We assume that this BChE staining is due to BChE activity anchored by PRiMA at the surface of glia cells as it was shown in CNS (Dobbertin *et al*., 2009). At higher magnifications though, a clear difference in the staining pattern of the heart hilum, the point of entry and exit of vessels, was observed between the genotypes. In WT, BChE^−/−^ and ColQ^−/−^ mice, intensely stained large fibre-like structures (black arrow in the Fig. 3) and small branching fibers (green arrows in the Fig. 3) were present. Small branching was not visible in AChE del E 5+6^−/−^ and PRiMA^−/−^ mice. Immunohistochemistry in transverse sections of the ventricle revealed co-localization of fluorescent signals for a specific anti-AChE antibody and antibody against the neuronal marker TUJ1 (Fig.3C).

**Figure 3.**
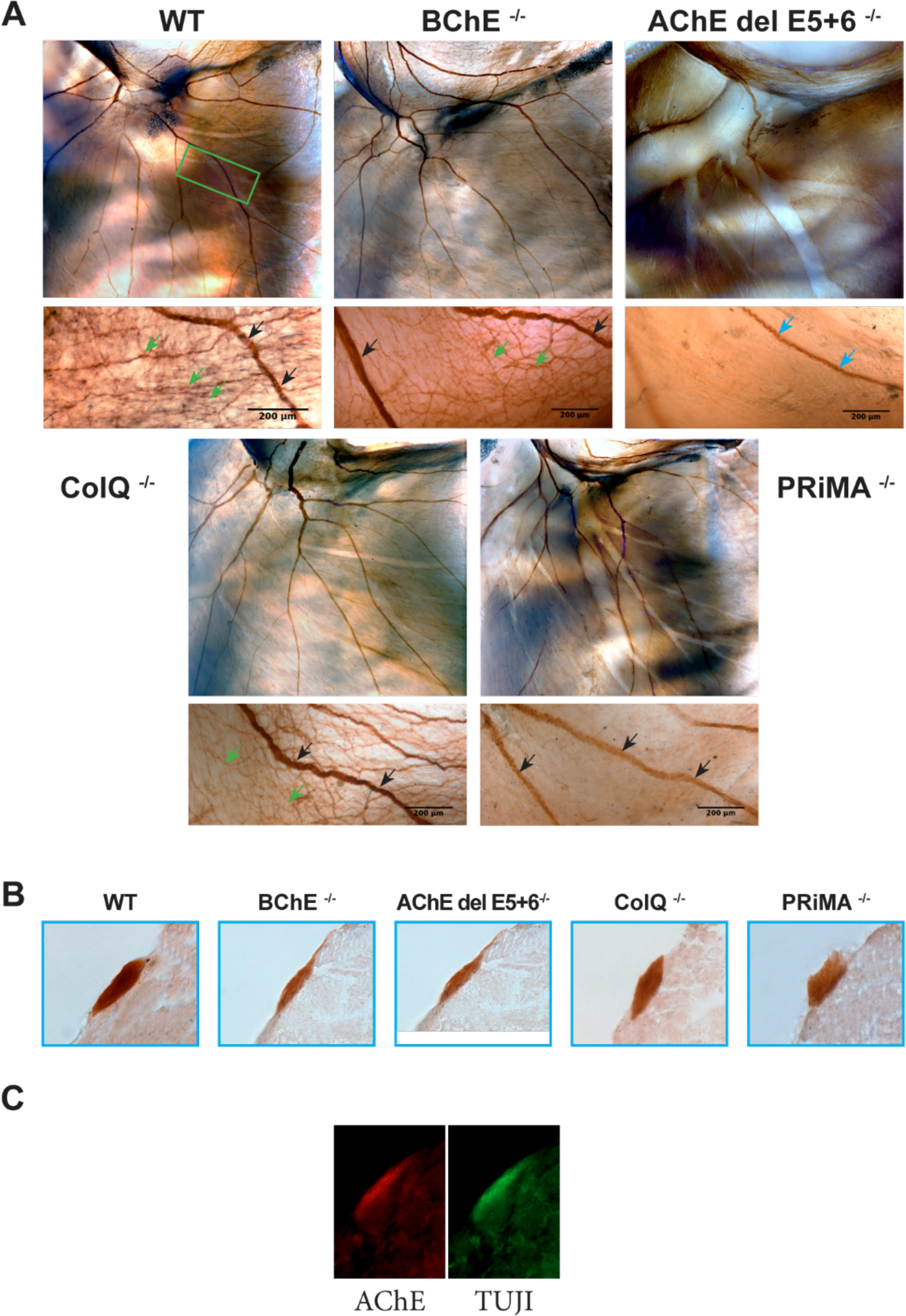
Localization of ChEs in the heart base. ***A,B*** Localization of ChE in ventricular epicardium of the whole-mounted heart (***A***) and in cryosections (***B***). ChE activity (marked by black arrows) was revealed for 3 hours with ATC as a substrate. Lower images depict the heart base at 4X magnification where strong ChE activity was detected in WT, BChE^−/−^, ColQ^−/−^, and PRiMA^−/−^ mice, but not in AChE del E 5+6^−/−^ mice, where only one coloured line was present (marked by blue arrows). The majority of staining in the heart base was formed by ColQ AChE and PRiMA AChE. The upper images depict hilum of the heart at 10X magnification, where the presence of AChE anchored by PRiMA in small branches of the heart hilum (marked by green arrows) were detected. This staining pattern is present in WT, BChE^−/−^, and ColQ^−/−^, but not in AChE del E 5+6^−/−^ and PRiMA^−/−^ mice. Images are representative of experiments that were repeated at least three times. **C** AChE co-localized with neuronal marker TUJ1 in immunohistochemistry.

Thus, our microscopy results confirm presence of both, PRiMA AChE and ColQ AChE, in epicardium of ventricles and moreover suggest that in contrast to the fiber-like structures, which seem to be formed almost equally by both PRiMA AChE and ColQ AChE, the small branching in the heart base was formed only by PRiMA AChE.

In ventricular myocardium, we did not detect any AChE activity in any of the studied genotypes, even after prolonged incubation. However, we observed strong diffuse BChE staining in WT compared to BChE^−/−^ mice, which was not affected by the heart perfusion process. Taken together with the results from biochemical analysis, BChE is clearly present in ventricular myocardium, presumably in endoplasmic reticulum where monomers are retained before assembly by PRAD into secretory tetramers.

### Heart physiology

Having established different localization of ChE molecular forms, we sought to evaluate the change of heart functions when ColQ or PRiMA anchors are inactivated or when BChE is absent.

### Left ventricular catheterization

We started with the direct recording of a pressure curve inside the left ventricle in anesthetized mice. Hemodynamic measurements revealed comparable basal values of heart rate in all studied mouse strains (WT: 380.4 ± 55.4 bpm, n = 8; PRiMA^−/−^: 360.4 ± 51.7 bpm, n = 6; BChE^−/−^: 382.1 ± 62.4 bpm, p = 0.6160, n = 7), that confirms preserved homeostatic regulation. Hemodynamic measurements were not possible to perform in ColQ^−/−^ and AChE Del E5+6^−/−^ due to the small size of the mice and fragility of the blood vessel. A dose-dependent positive chronotropic effect of the β_1_-adrenergic agonist dobutamine was observed in all studied mouse strains. Nevertheless, the magnitude of this effect depended on the genotype (Table 1). While the heart rate increase was similar in WT and BChE^−/−^ mice, the response of PRiMA^−/−^ hearts to dobutamine at concentrations ≥ 50 µg/kg (p = 0.0030 for 50 µg/kg, p = 0.0310 for 100 µg/kg, p = 0.0396 for 166.8 µg/kg) was significantly higher compared to WT (Table 1).

**Table 1:** Heart rate increase in WT and mutant mice during dobutamine application.

| Dobutamine dose<br>[μg/kg] | WT<br>[%] ± SD | PRiMA<sup>-/-</sup><br>[%] ± SD | BChE<sup>-/-</sup><br>[%] ± SD |
| --- | --- | --- | --- |
| 0.0 | 100 | 100 | 100 |
| 16.7 | 110.1 ± 4.0 | 115.2 ± 5.1 | 116.9 ± 10.5 |
| 50.0 | 116.0 ± 2.2 | **128.3 ± 5.2 (\*\*)** | 124.1 ± 16.5 |
| 100.0 | 128.2 ± 4.2 | **139.7 ± 7.6 (\*)** | 133.3 ± 19.2 |
| 166.8 | 133.7 ± 4.3 | **145.7 ± 8.1 (\*)** | 141.5 ± 17.1 |

Values are reported as mean ± SD; Heart rate values of PRiMA<sup>-/-</sup> against WT values (p = 0.0030 for 50 μg/kg, p = 0.0310 for 100 μg/kg, p = 0.0396 for 166.8 μg/kg); WT, n = 6; PRiMA<sup>-/-</sup>, n = 6; BChE<sup>-/-</sup>, n = 6.

Basal hemodynamic parameters LVSP, LVDP, dP/dt_max_ and dP/dt_min_ of mutant mice were similar to WT. After administration of dobutamine, PRiMA^−/−^ and WT mice responded similarly. However, LVSP in BChE^−/−^ was less than WT mice in response to dobutamine (Table 2, p = 0.0358 for 16,7 µg/kg, p = 0.0320 for 50 µg/kg, p = 0.0032 for 100 µg/kg, p = 0.0009 for 166.8 µg/kg). The same was observed for the rate of contraction (dP/dt_max_) (Table 2, p = 0.0052 for 16,7 µg/kg, p = 0.0325 for 50 µg/kg, p = 0.0068 for 100 µg/kg, p < 0,0001 for 166.8 µg/kg) but not for the rate of relaxation (−dP/dt_min_) where significant change was observed solely for the highest dobutamine dose (Table 2, p < 0,0001).

**Table 2:** LVSP, dP/dt_max_ and dP/dt_min_ values in WT and mutant mice during dobutamine application.

| Dobutamine dose<br>[μg/kg] | WT<br>[%] ± SD |  | PRiMA<sup>-/-</sup><br>[%] ± SD |  | BChE<sup>-/-</sup><br>[%] ± SD |  |
| --- | --- | --- | --- | --- | --- | --- |
|  | LVSP | dP/dt<sub>max</sub> | LVSP | dP/dt<sub>max</sub> | LVSP | dP/dt<sub>max</sub> |
|  |  | -dP/dt<sub>min</sub> |  | -dP/dt<sub>min</sub> |  | -dP/dt<sub>min</sub> |
| 0.0 | 100 | 100 | 100 | 100 | 100 | 100 |
|  |  | 100 |  | 100 |  | 100 |
| 16.7 | 112.2 ± 3.0 | 130.7 ± 6.3 | 110.2 ± 2.9 | 140.0 ± 10.1 | 107.0 ± 1.9 (*) | 112.9 ± 9.7 (**) |
|  |  | 121.3 ± 9.0 |  | 113.9 ± 4.0 |  | 114.1 ± 7.0 |
| 50.0 | 114.1 ± 3.9 | 134.8 ± 8.8 | 112.1 ± 4.5 | 151.6 ± 10.1 (*) | 108.8 ± 2.0 (*) | 119.9 ± 9.7 (*) |
|  |  | 123.0 ± 13.5 |  | 121.2 ± 7.4 |  | 117.1 ± 6.5 |
| 100.0 | 120.9 ± 3.9 | 149.0 ± 9.0 | 119.0 ± 7.3 | 161.9 ± 10.5 | 113.8 ± 2.4 (**) | 131.6 ± 11.0 (**) |
|  |  | 133.2 ± 13.3 |  | 129.7 ± 12.2 |  | 124.7 ± 8.1 |
| 166.8 | 126.8 ± 2.1 | 174.0 ± 4.6 | 125.3 ± 8.3 | 167.8 ± 12.8 | 118.8 ± 2.1 (***) | 138.8 ± 8.8 (****) |
|  |  | 159.5 ± 11.3 |  | 138.4 ± 15.8 (***) |  | 131.1 ± 7.8 (****) |

Values are reported as mean ± SD; LVSP of BChE<sup>-/-</sup> against WT values (p = 0.0358 for 16,7 µg/kg, p = 0.0320 for 50 µg/kg, p = 0.0032 for 100 µg/kg, p = 0.0009 for 166.8 µg/kg); dP/dt<sub>max</sub> of BChE<sup>-/-</sup> against WT values (p = 0.0052 for 16,7 µg/kg, p = 0.0325 for 50 µg/kg, p = 0.0068 for 100 µg/kg, p < 0,0001 for 166.8 µg/kg); dP/dt<sub>max</sub> of PRiMA<sup>-/-</sup> against WT values (p = 0.0100 for 50 µg/kg); dP/dt<sub>min</sub> of BChE<sup>-/-</sup> against WT values (p < 0,0001 for 166.8 µg/kg); dP/dt<sub>min</sub> of PRiMA<sup>-/-</sup> against WT values (p 0.0003 for 166.8 µg/kg); WT, n = 6; PRiMA<sup>-/-</sup>, n = 6; BChE<sup>-/-</sup>, n = 6.

Thus, lack of molecular forms of ChE does not influence basal hemodynamic parameters. Nevertheless, knock out of PRiMA AChE or BChE leads to different responses to dobutamine, with a higher heart rate increase or a lower LVSP, respectively.

### Isolated perfused heart assay

To study heart physiology without interference from *in vivo* regulatory and compensatory mechanisms, isolated hearts were mounted on a Langendorff’s perfusion system. Functional parameters were measured in spontaneously beating or paced hearts in basal conditions and under cholinergic stimulation.

### Spontaneously beating hearts

In basal conditions, there was no significant difference in heart rate between the mouse strains (WT: 296.0 ± 79.4 bpm, n = 10; PRiMA^−/−^: 298.7 ± 69.1 bpm, n = 9; AChE del E5+6^−/−^: 306.1 ± 135 bmp, n = 16; BChE^−/−^: 246.0 ± 92.0 bpm, n = 9), with exception of ColQ^−/−^ hearts where significantly higher values (416.2 ± 52.6 bpm, n = 18, p = 0.0057) were observed when compared to WT.

Neostigmine, a ChE inhibitor, was used to block the degradation of ACh that is spontaneously secreted. In WT mice, we observed a gradual decrease in heart rate within the first 10 min of neostigmine application, which then stabilized for the remainder of the 25 min (Fig. 4 left panel). In a second set of experiments, the muscarinic receptor antagonist atropine was added after 10 min application of neostigmine, and the heart rate recovered after 2 min (Fig. 4 right panel). AChE del E 5+6^−/−^ hearts did not respond to neostigmine (p = 0.013 (5 min), p = 0.0147 (10 min), p = 0.0094 (15 min), p = 0.0106 (20 min), p = 0.0075 (25min) against WT values) and atropine had no effect on heart rate. This result established that in this experimental design only AChE and not BChE hydrolysed spontaneously released ACh. We anticipated a similar resistance in hearts from PRiMA^−/−^ mice as PRiMA is the anchor of AChE to the plasma membrane of neuron. Nevertheless, we found a decrease of heart rate and recovery after atropine comparable to WT. This suggests that ColQ AChE hydrolyses ACh in the absence of AChE anchored to the membrane of neurons. Interestingly, a two-fold decrease in heart rate was observed during neostigmine application in BChE^−/−^ hearts as compared to WT (p = 0.023 (5 min), p = 0.0376 (10 min), p = 0.0193 (20 min)) (Fig. 4 left panel). Also, in contrast to other genotypes, the effect of neostigmine was reversed gradually after atropine application (1 µM) (Fig. 4 right panel). The nicotinic receptor antagonist hexamethonium (100 µM) had no effect on heart rate in neostigmine treated hearts.

**Fig. 4.**
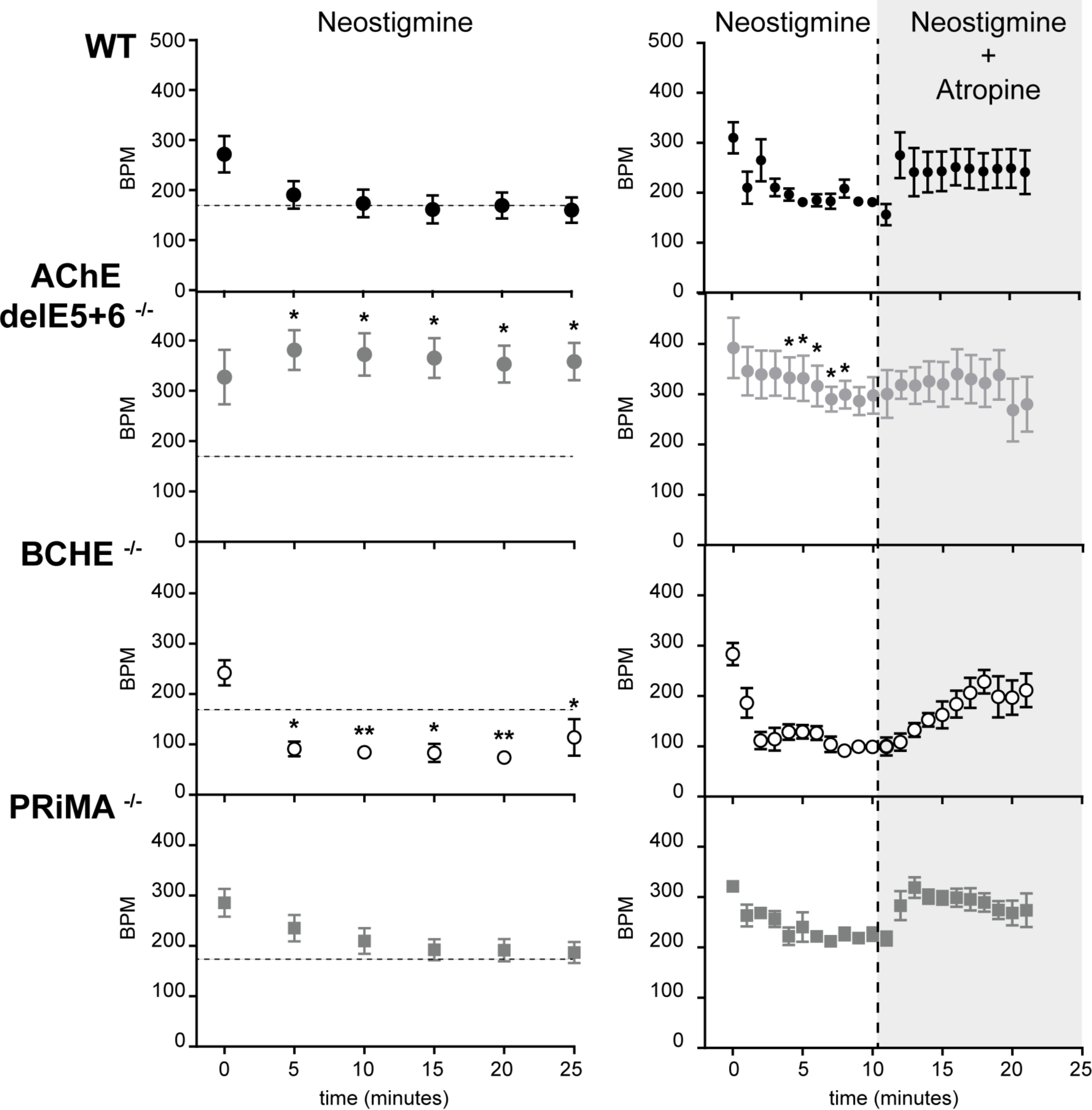
Heart rate changes upon neostigmine administration. ***Left panel*** Comparison of the decrease in heart rate during application of 3 μM neostigmine for 25 minutes between WT mice and mutant mice. AChE del E 5+6^−/−^ mice did not respond to neostigmine application (p = 0.013 (5 min), p = 0.0147 (10 min), p = 0.0094 (15 min), p = 0.0106 (20 min), p = 0.0075 (25 min) against WT values), BChE^−/−^ mice showed a much more pronounced decrease in heart rate as compared to WT (p = 0.023 (5 min), p = 0.0376 (10 min), p = 0.0193 (20 min)), and PRiMA^−/−^ mice were comparable to WT. WT: n = 8, AChE del E 5+6^−/−^: n = 5, BChE^−/−^: n = 10, PRiMA^−/−^: n = 9. ***Right panel*** Comparison of the decrease in heart rate during application of 3 μM neostigmine and subsequent muscarinic receptor blockade with 1 μM atropine in WT and AChE del E 5+6^−/−^ mice, BChE^−/−^ mice, PRiMA^−/−^ mice. WT: n = 4, AChE del E 5+6^−/−^: n = 4, BChE^−/−^: n = 6, and PriMA^−/−^: n = 4. Values are presented as mean ± SEM.

Thus, the reduction of heart rate due to ChE inhibitor neostigmine results presumably from the canonical activation of muscarinic M2 receptors on cardiomyocytes by ACh released by the parasympathetic neurons. ACh is controlled by AChE, anchored by ColQ or PRiMA. In isolated heart of BChE^−/−^, the accentuated decrease of heart rate after neostigmine could depend on adaptation of heart from the total absence of BChE.

The cholinergic system counteracts the beta-adrenergic action, we thus in the next step stimulated β-adrenergic receptors with isoproterenol (ISO). As anticipate, we observed a concentration-dependent increase in heart rate in all genotypes (Fig. 5). However, the maximal percent increase in heart rate induced by ISO was less in ColQ^−/−^ hearts due to higher basal heart rate (Fig. 5, p = 0.0048 (KREBS), p = 0.0161 (0.1nM ISO), p = 0.0231 (0.3nM ISO), p = 0.0391 (1nM ISO) against WT values).

**Fig. 5.**
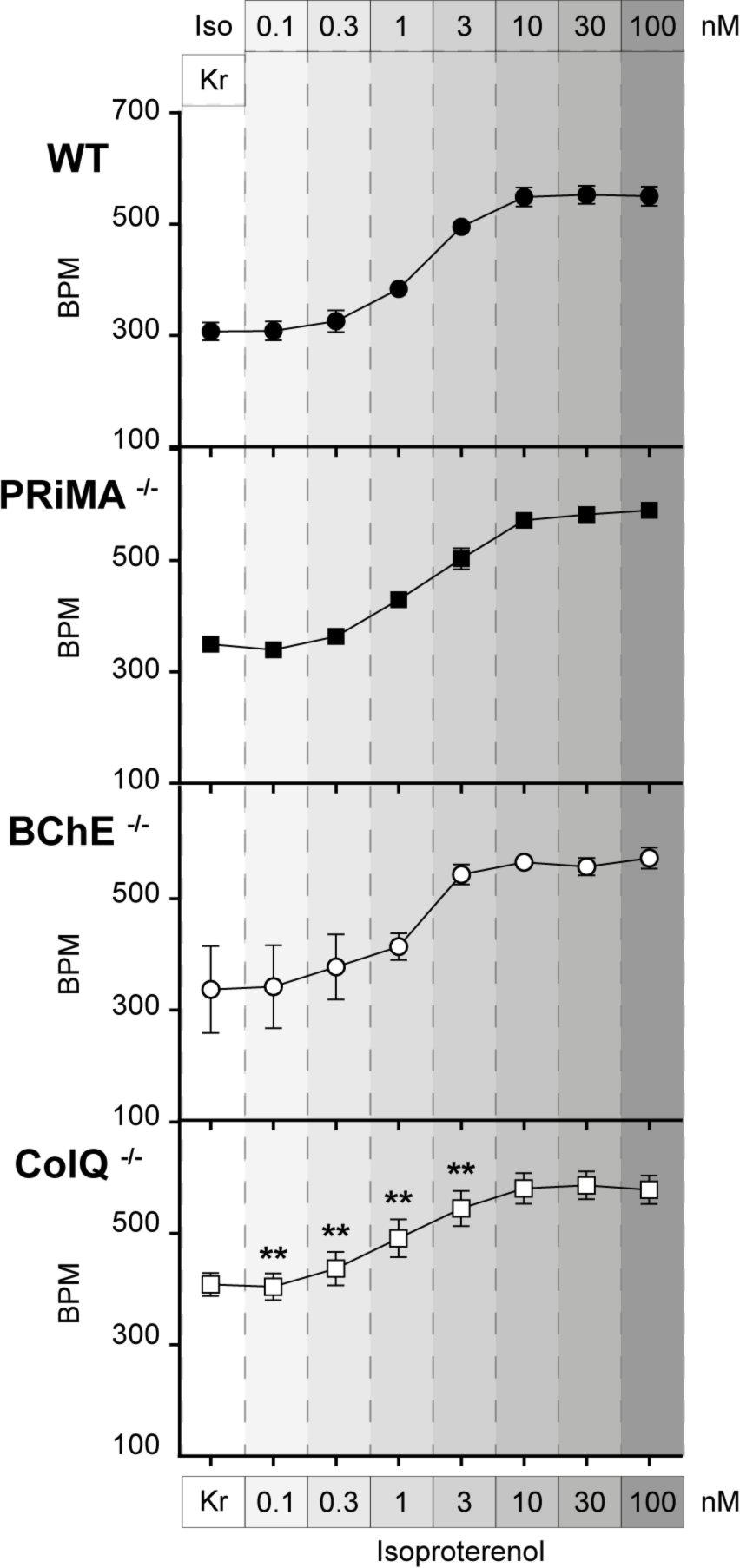
Comparison of heart rate during isoproterenol (ISO) administration. The graphs present the mean of heart rates, in basal condition (Kr) and after 7 min incubation in medium containing increasing concentration of isoproterenol (ISO). Dose-effect relationship was similar in WT mice (n = 8), BChE^−/−^ mice (n = 3) and PRiMA^−/−^ mice (n = 7). The maximal percentage increase in heart rate induced by ISO was less in hearts of ColQ^−/−^ mice (n = 8) due to higher basal heart rate (p = 0.0048 (KREBS), p = 0.0161 (0.1nM ISO), p = 0.0231 (0.3nM ISO), p = 0.0391 (1nM ISO) against WT values). Values are reported as mean ± SEM

We next tested for the capacity of ACh to antagonize the β-adrenergic response to ISO. In all studied strains, ACh decreased heart rate when added on top of ISO (100 nM, Fig. 6 *left*). Interestingly, BChE^−/−^ hearts responded differently to the highest ACh concentration (10 µM, p = 0.0255), at which heart rate was significantly higher than heart rate recorded for WT hearts, i.e. the heart appears less sensitive to ACh.

**Fig. 6.**
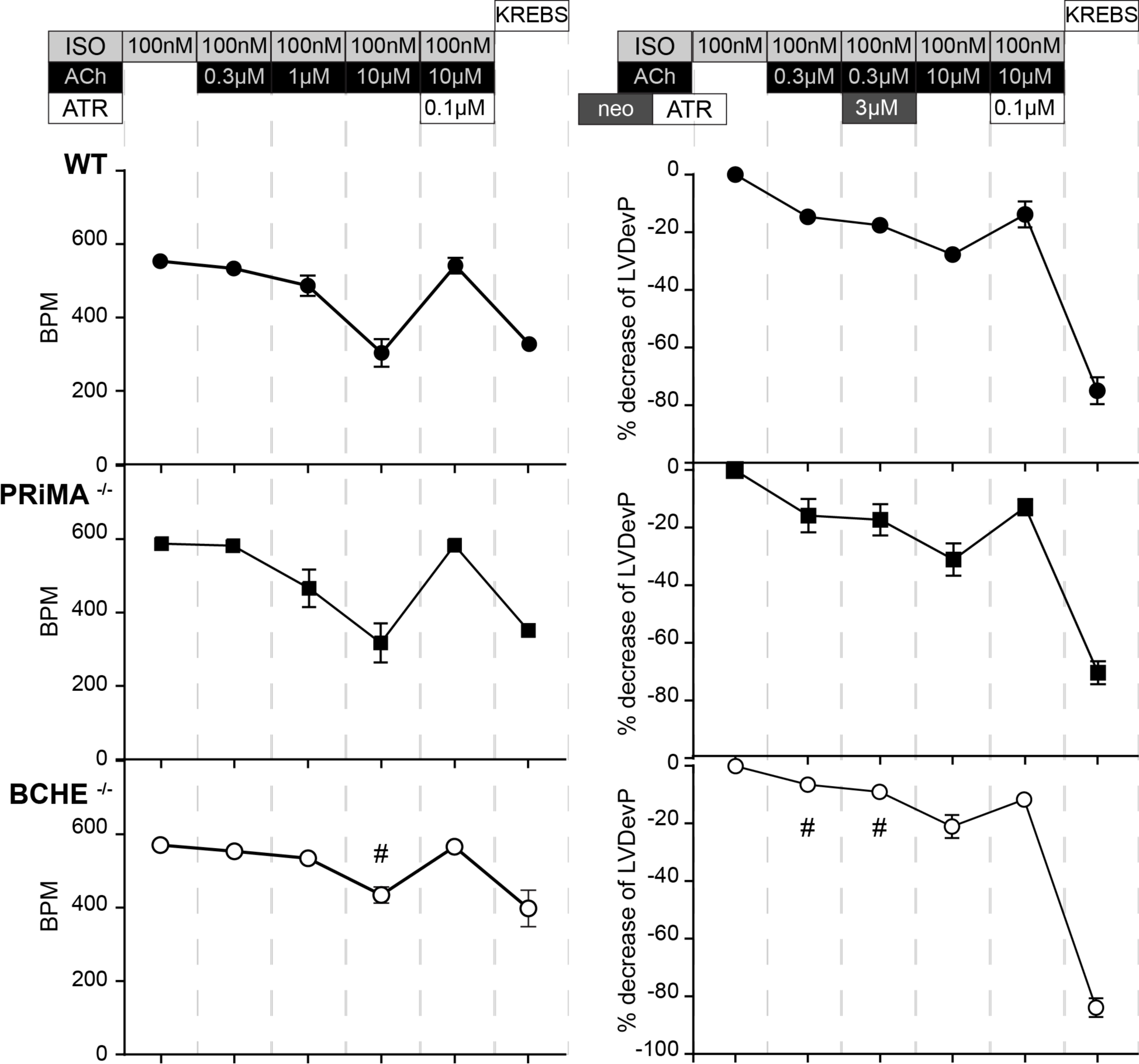
Effect of acetylcholine (ACh) application on heart rate of isoproterenol (ISO) pretreated hearts. ***Left panel*** The graphs present the mean values of heart rate recorded in spontaneously beating Langendorff’s isolated hearts. Hearts were treated with isoproterenol (ISO 100nM) and then incubated with different concentrations of acetylcholine (ACh) and the effect was reversed subsequently by application of atropine (ATR). Hearts of WT mice (n=8) and PRiMA^−/−^ mice (n = 7) responded similarly to the ACh applications at all doses. Hearts from BChE^−/−^ mice (n = 3) responded differently to the 10 µM ACh concentration (p = 0.0255). ***Right panel*** The graphs present the variation of Left Ventricular Developed Pressure (LVDevP) obtained from Langendorff’s perfused hearts paced to 650 bpm pretreated with isoproterenol (ISO 100nM). Action of ISO was reversed with ACh (0.3µM) with or without application of neostigmine (NEO, 3µM). Hearts of WT mice (n=3) and PRiMA−/− mice (n=6) responded similarly. In contrast, hearts from BChE^−/−^ mice responded less to the cholinergic stimulation: p = 0.0094 (0.3 µM ACh), p = 0.0151 (3 µM NEO) against WT values. At higher concentration of ACh (10µM), the pressure decreased in all genotypes and was reversed by atropine. After washing with Krebs the pressure decreased dramatically. Values are reported as mean ± SEM.

### Paced hearts

When isolated hearts were paced at 650 bpm, basal isovolumic contraction showed similar values in all studied genotypes for Left Ventricular Developed Pressure (LVDevP) (WT: 43.9 ± 9.8 mmHg, n = 5, BChE^−/−^: 41.3 ± 6.8 mmHg, n = 6, AChE del E 5+6^−/−^: 41.6 ± 7.6 mmHg, n = 7, p = 0.8388), for End-Diastolic Pressure EDP (WT: 36.6 ± 12.8 mmHg, n = 5, BChE^−/−^: 37.7 ± 15.6 mmHg, n = 6, AChE del E 5+6^−/−^: 35.7 ± 12.7 mmHg, n = 7, p = 0.9654), for dP/dtmax (WT: 2080.8 ± 730.8 mmHg/s, n = 5, BChE^−/−^: 2010.1 ± 640.6 mmHg/s, n = 6, AChE del E 5+6^−/−^: 1751.3 ± 358.2 mmHg/s, n = 7, p = 0.5414), and for dP/dtmin (WT: −1281.7 ± 174.3 mmHg/s; n=5, BChE^−/−^: −1289.9 ± 219.3 mmHg/s; n=6, AChE del E 5+6^−/−^: −1201.7 ± 191.2 mmHg/s; n=7, p = 0.6578). Inhibition of ChE by neostigmine had no effect on all measured parameters. β-adrenergic stimulation with ISO led to a dose dependent increase of LVDevP, dP/dtmax and -dP/dtmin and decrease of EDP in all studied strains. Decrease of LVDevP in response to cholinergic stimulation by an exogenous ACh was comparable in all three genotypes. However, BChE^−/−^ hearts responded by a significantly lower decrease of LVDevP during cholinergic stimulation of the isoproterenol-treated hearts (Fig. 6*B*, p = 0.0094 (0.3 µM ACh), p = 0.0151 (3 µM NEO) against WT values).

## Discussion

Since inhibition of ChEs is one of the promising cholinergic therapeutic approaches in cardiovascular diseases (Roy *et al*., 2015), there is an urgent need to characterize these enzymes in the heart. Here, we leveraged genetically modified mouse lines to establish the molecular origins, distribution, and physiological contribution of AChE and BChE in the heart. We confirmed previous reports of higher BChE than AChE activity in the whole mouse heart (Gómez *et al*., 1999; Li *et al*., 2000). However, when studied in separated heart regions, we found a unique ChE distribution.

### AChE predominates in heart atria, and colocalizes with neurons

Similarly to other species (Sinha *et al*., 1976; Nyquist-Battie & Trans-Saltzmann, 1989), the highest AChE activity was detected in the mouse atria. It was present in anchored forms, with PRiMA AChE slightly exceeding ColQ AChE. We observed that areas with lower AChE activity, i.e., the right ventricle, left ventricle and septum, showed slightly higher ColQ AChE content.

The precipitate due to AChE activity was absent in atria of AChE delE5+6 mice in which AChE cannot interact with PRiMA or ColQ and in PRiMA−/− atria. Thus, AChE staining in atria was due to AChE anchored by PRiMA. Despite significant quantity of ColQ AChE detected in atrial protein extracts, we were unable to visualize ColQ AChE in the studied layers of the PRiMA^−/−^ cardiac atria. We assume that in this heart chamber, ColQ AChE is not concentrated in one structure but is dispersed within the tissue and thus is undetectable. A similar discrepancy between biochemical and microscopic analysis was observed in skeletal muscles, where quantities of PRiMA AChE and ColQ AChE were similar, but only ColQ AChE was visible at the neuromuscular junction. PRiMA AChE was extracted from muscle parts that do not contain neuromuscular junction. It was diffusely distributed along the muscle, in the extrajunctional regions (Bernard *et al*., 2011). Nevertheless, we cannot exclude that prolonged tissue handling in microscopy may have resulted in faster degradation of ColQ AChE. This assumption is, however, weakend by the fact that microscopic ColQ AChE detection problems were specific to the atria, since in other heart chambers and compartments, ColQ AChE protein and activity were detected.

In the heart base, we visualized both ColQ AChE and PRiMA AChE. Dense AChE staining that was present in PRiMA^−/−^ or ColQ ^−/−^ hearts but missing in AChE delE5+6^−/−^ mice extended through the epicardia to the atria but was absent in myocardium. Apart from strongly coloured fibre-like structures, we also observed subtle branches present in ColQ^−/−^ hearts but not in PRiMA^−/−^ or AChE delE5+6^−/−^ hearts thus consisting of PRiMA AChE.

AChE has commonly been used to identify neurons in cardiac tissue, both the ganglia and the nerves (Rysevaite *et al*., 2011; Pauza *et al*., 2013). At ultrastructural level, AChE was observed at the surface of the neurons but not in the cardiomyocyte (Silva *et al*., 1993). Indeed, the canonical function of PRiMA is to organize AChE into tetramers and target them to the surface of the cholinergic neurons, as shown in brain (Perrier *et al*., 2002, 2003; Dobbertin *et al*., 2009). PRiMA AChE was also shown to be produced by motoneurons (Bernard *et al*., 2011). Our results of co-localization with the neuronal marker TUJ1 confirm neuronal origin of AChE. Nevertheless, ColQ attaches AChE to the basal lamina and thus AChE should not be strictly used as a neuronal marker.

### AChE participates in heart rate regulation

In the isolated heart, release of ACh is not triggered by an external signal i.e. electrical stimulation. Cholinergic stimulation may be achieved by a direct or indirect agonistic effect on M2 muscarinic receptors. Both the direct muscarinic receptor agonist, ACh, and indirect agonist, nonselective ChE inhibitor neostigmine, decreased heart rate of the isolated heart. The lack of bradycardic effect on AChE delE5+6 ^−/−^ heart confirmed that the level of ACh is controlled by AChE. Moreover, a comparable heart rate decrease in WT, PRiMA^−/−^ and ColQ ^−/−^ (not shown) hearts proves the participation of both cardiac AChE forms, PRiMA AChE and ColQ AChE.

Systemic administration of the β_1_-adrenergic receptor agonist, dobutamine, to PRiMA^−/−^ led to a higher heart rate increase compared to WT mice. Nevertheless, this discrepancy upon adrenergic stimulation was not observed in Langendorff perfused hearts, suggesting that the adrenergic effect on heart rate in PRiMA^−/−^ mice was facilitated by non-cardiac tissues. We postulate that the CNS might be involved, as the tissue is rich in PRiMA AChE (Dobbertin *et al*., 2009) and dobutamine has some α-adrenergic properties (Ruffolo *et al*., 1981) that are important in central heart rate regulation (Haeusler, 1975).

### BChE predominates in the ventricles as intracellular monomer, precursor of secreted tetramer

We observed BChE activity distribution within heart chambers/regions to be opposite to that of AChE. The lowest BChE activity, although undetectable by microscopy, was recorded in the atria. In ventricles, BChE activity was diffusely distributed, likely due to intracellularly localized monomer. Additionally, we consistently observed one strongly colored fibre-like structure in the base of the heart. The precise localization suggests the presence of anchored forms of BChE, probably PRiMA BChE. Gómez et al. (1999) showed small amounts (3%) of this form in the mouse heart (Gómez et al., 1999). In our experimental conditions, we were not able to detect this low level of PRiMA BChE activity most likely because we solubilized all the proteins in a single extraction whereas Gomez et al. used sequential extractions. We found mainly soluble BChE monomer, which is a precursor of tetramer organized by proline rich peptide and secreted (Blong Rm *et al*., 1997). BChE monomers are retained in the endoplasmic reticulum and are not in a position to hydrolyze ACh (Schopfer Lm & Lockridge O, 2016).

### Adaptation of heart function in absence of BChE

Despite BChE not being quantitively anchored to the plasma membrane or deposited in the basal lamina, the heart rate of BChE^−/−^ isolated hearts decreased more than that of WT or PRiMA^−/−^ hearts when perfused with 3µ neostigmine. As discussed above, neostigmine action in heart depends on AChE inhibition thus changes observed in BChE^−/−^ suggest adaption to the chronic absence of BChE that may be due to increased release of ACh in non-stimulated heart, and thus increased M2 muscarinic receptors stimulation and/or by adaptation at the level of M2 muscarinic receptor signaling pathway.

To reveal the character of the adaption, two points should be considered. First, there was an increased response to the indirect agonist on muscarinic receptors neostigmine. Neostigmine inhibits degradation of ACh produced by heart and thus increases its extracellular levels. Nevertheless, an application of ACh that directly stimulates the receptors had not such effect but in contrary led to lesser action on heart rate under adrenergic stimulation in BChE^−/−^ mice than WT mice. We therefore conclude that heart of BChE^−/−^mice release more ACh than WT, presumably to adapt heart rate. Second point is different dynamics of atropine muscarinic-receptor-blocking action in neostigmine treated hearts and different response of isoproterenol-stimulated hearts to ACh which suggest adaptations at the level of muscarinic receptors. Receptor adaptation has also been shown in brains of mutant mice (Volpicelli-Daley et al., 2003; Hrabovska et al., 2010; Farar et al., 2012) in which an approximately 300-fold increase of ACh was detected due to the reduction of ChE (Hartmann et al., 2007; Farar et al., 2013). Thus, hearts of BChE^−/−^ mice seem to adapt heart rate by changes in both, ACh release and muscarinic receptor signaling.

A possible regulatory function of BChE was also observed during inotropic studies as a slower increase of LVSP and dP/dt_max_ was observed during β_1_-adrenergic stimulation in BChE^−/−^, compared to WT mice. Also, addition of exogenous ACh to paced hearts with stimulated adrenergic system caused a slower decrease of LVDevP in BChE^−/−^ than in WT mice. Thus, the absence of BChE may lead to higher levels of ACh in the ventricles, which antagonizes the positive inotropic effect of β-adrenergic stimulation.

Two important questions sustain from our results. First, what is the origin of released ACh and second, how may BChE although abundant in ventricle but not immediately available to hydrolyze ACh affect the release of ACh. A positive effect of cholinergic activation by direct vagal stimulation or ChE inhibition has been confirmed in animal models as well as humans, and a role of both neuronal and non-neuronal systems has been documented (Roy *et al*., 2015). It is well known that the cholinergic system provides negative feedback to control the chronotropic responses to sympathetic drive. Interestingly, genetic inactivation of ACh synthesis in cardiomyocytes leads to a delay in heart rate recovery after exercise in mice showing the importance of cardiomyocyte ACh in this process (Roy et al., 2013). Our observation that hearts lacking BChE were less sensitive to ACh as a 10 µM concentration was unable to fully block the β-adrenergic effect on heart rate revealed a role of BChE in heart rate recovery under sympathetic stimulation. This further strengthens our suggestion that BChE, as part of the non-neuronal cholinergic system in heart, is responsible for the hydrolysis of cardiomyocyte-produced ACh. Another open question is which cell could be the target of nonneuronal ACh release by cardiomyocyte of ventricle to regulate heart rate. In lung, nonneuronal ACh is released by epithelial cells named tuft or guard cells. ACh released by non-neuronal cell activates vagal afferent neurons and control the respiration (Krasteva *et al*., 2011). By analogy with the role of ACh in lung, we propose that ACh from the cardiomyocytes may activate vagal afferent neurons and control heart rate. It was shown that specific stimulation of afferent vagal nerve stops heart to beat and stop the ventilation of mice (Chang *et al*., 2015). It is thus tempting to propose that vesicles of ACh released by cardiomyocyte activate afferent vagal nerve that secondary reduce heart rate through the canonical parasympathetic control. Adaptations of BChE^−/−^ mice may reveal indirectly the synergic action of ACh in the control of heart rate demonstrated by the genetic deletion of ACh in cardiomyocyte.

### Limits and general considerations

Based on the overall analysis in mice, we can propose that AChE anchored to the membrane of the neurons by PRiMA and to the extracellular matrix by ColQ hydrolyses neuronal ACh while BChE hydrolyses cardiomyocyte ACh, at least when BChE is in the extracellular space. This global distribution of ChE varies with the species, in rat, BChE is anchored by ColQ in ventricle and thus part of the enzymes is anchored in the extracellular matrix (Nyquist-Battie & Trans-Saltzmann, 1989; Krejci *et al*., 1997).

Although we evaluated physiological parameters of heart, we did not directly address the role of AChE in the control of M2 muscarinic receptors activity that control G protein-activated inwardly rectifying K1 channels (Sakmann *et al*., 1983). Cardiomyocytes are highly sensitive to ACh, and the termination of activation of M2 muscarinic receptors does not depend on ACh hydrolysis, thus the function of AChE that we reveal with neostigmine is rather the distance of diffusion of ACh from a cholinergic bouton to cardiomyocytes than the termination of the action of ACh in synapse. Even in the cholinergic synapse, the absence or inhibition of AChE changes the activation of the post-synaptic receptors but ACh spillover activates more extra-synaptic receptors (Lamotte d’Incamps *et al*., 2012). This simple view is complicated by the fact that the peripheric neuronal circuits that regulate heart rate are complex (Rajendran *et al*., 2019). In this complex net, AChE is localized in cholinergic and adrenergic neurons. Recent studies revealed that neuronal ACh but not cardiomyocyte ACh contributes to an autonomic imbalance contract by pyridostigmine a non-specific ChEI (Guimarães *et al*., 2022).

Pyridostigmine is a non-specific ChEI that was broadly used for the treatment of myasthenic syndrome, when the density of functional muscle nicotinic receptors is low at the neuromuscular junction (Lorenzoni *et al*., 2020), or as protector against nerve agents. In both cases, the side effects are numerous presumably because ACh is used not only in skeletal muscle but also in several tissues in which BChE is more abundant than AChE. Indeed, 30% of American soldiers who took pyridostigmine during Iraq war suffered of Gulf syndrome (Golomb, 2008). A part of this heterogeneity has a genetic origin (Steele *et al*., 2015). To follow the rational of proposed therapeutic potential of cholinergic activation by ChEI in heart failure (Roy et al., 2015), ours results reveal a complex coordination of ChE in mouse heart that must be evaluated in human heart, knowing that the distribution of AChE and BChE is different in blood of mouse and human. It is thus conceivable that the efficacy of the ChEIs depends not only on the inhibitor used but also on the polymorphism of BChE in human.

## Data availability statement

All data contained in this study are available from the authors upon reasonable request.

## Competing interests

The authors declare that they have no competing interests.

## Author contributor statement

The hypothesis was formed, and the study was designed by [A.H.], [E.K.] and [R.F.]. Material preparation, data collection and analysis were performed by [D.D.], [M.K.] and [T.H.]. The first draft of the manuscript was written by [A.H.] and [D.D.] and all authors commented on previous versions of the manuscript. All authors read and approved the final manuscript.

## Funding

This work was supported by The Slovak Research and Development Agency [grant number APVV-22-0541 to A.H.]; The Ministry of Education, Science, Research and Sport of the Slovak Republic [grant numbers VEGA 1/0815/21 to A.H, VEGA 1/0283/22 to D.D], by research grants (21008; 23138) from AFM Telethon to EK.

## Acknowledgements

We would like to thank Dr. Evan David Paul for insightful comments and proofreading, Roman Dinga for IT and graphical support and prof. Palmer Taylor for providing us with primary anti-AChE antibodies.

## Notes

### Competing Interest Statement

The authors have declared no competing interest.

